# Oligomeric polymorphism of HIV-1 Vpu protein in lipid environment and in solution

**DOI:** 10.1101/2022.08.26.505453

**Authors:** Saman Majeed, Oluwatosin Adetuyi, Md Majharul Islam, Bo Zhao, Elka R. Georgieva

## Abstract

The HIV-1 encoded protein Vpu forms an oligomeric ion channel/pore in membranes and interacts with multiple host proteins to support virus lifecycle. However, Vpu molecular mechanisms are currently not well understood. The structures of full-length Vpu in its monomeric and oligomeric forms are unknown, although both the monomer and oligomer are deemed important. Here, we report on the diversity of Vpu oligomeric structures and how the environment affects the Vpu oligomer formation. We produced a uniquely designed MBP-Vpu chimera protein in *E. coli* in soluble form. We subjected this protein to analytical size exclusion chromatography (SEC) and negative staining electron microscopy (nsEM). Strikingly, we found that MBP-Vpu forms stable oligomers in solution, presumably driven by Vpu transmembrane domain self-association. Our coarse modeling suggests that these oligomers are pentamers, in agreement with the pentameric membrane-bound Vpu. To the best of our knowledge, this is the first observation of Vpu self-association out of lipid membrane environment. We further found that MBP-Vpu oligomer stability decreases when the protein was reconstituted in lipid membrane mimetics, such as β-DDM, and mixtures of lyso PC/PG or DHPC/DHPG—In these cases significant oligomer heterogeneity was observed with oligomeric order lesser than that of MBP-Vpu oligomer in solution, but larger oligomers were observed as well. Importantly, we found that in lyso PC/PG, above certain protein concentration, MBP-Vpu forms linear array-like structures, which is also novel. Thus, our studies provide unique information about Vpu protein quaternary organization by capturing multiple Vpu oligomeric structures, which we believe are physiologically relevant.

## INTRODUCTION

Human immunodeficiency virus 1 (HIV-1) belongs to the family of *Retroviridae* and is the causative agent of HIV/AIDS.(1) Worldwide, each year more than 30 million people carry the virus and close to 700,000 of them die due to virus-related causes.(2) Regrettably, the resources to curb the virus and cure the HIV-caused health complications are still limited. Depending on the origin and spread within human population, the virus was divided into M, N, O, and P subtypes, with M subtype being globally distributed and the major HIV-1 pandemic factor.(3) Detailed knowledge about the molecular mechanisms the virus uses in the infected cells is needed to comprehensively understand HIV-1 pathogenesis and develop means to combat the virus. In this regard, one common for all HIV-1 subtypes and at the same time outstanding HIV-1 protein is Vpu (viral protein U)—Among other important proteins, all HIV-1 subtypes encode the accessory protein Vpu; but, strikingly, it was found that this protein is most active in the pandemic-causing subtype M, suggesting that Vpu has an important role in HIV-1 lifecycle and infectivity.(1,3,4) Vpu is expressed in the infected cells and is located in the membranes of trans-Golgi network, endoplasmic reticulum (ER) and plasma membrane.(4,5) It is a small protein made of about 81 amino acids; it is divided into an N-terminal domain with conserved hydrophobic helical region (Helix 1, transmembrane domain [TMD]), which resides in and traverses the cellular lipid membranes, and a C-terminal domain containing two helical regions, Helix 2 and Helix 3, with Helix 2 possibly being associated with membrane surfaces (Figure 1).(1,6-8)

**Figure 1.**
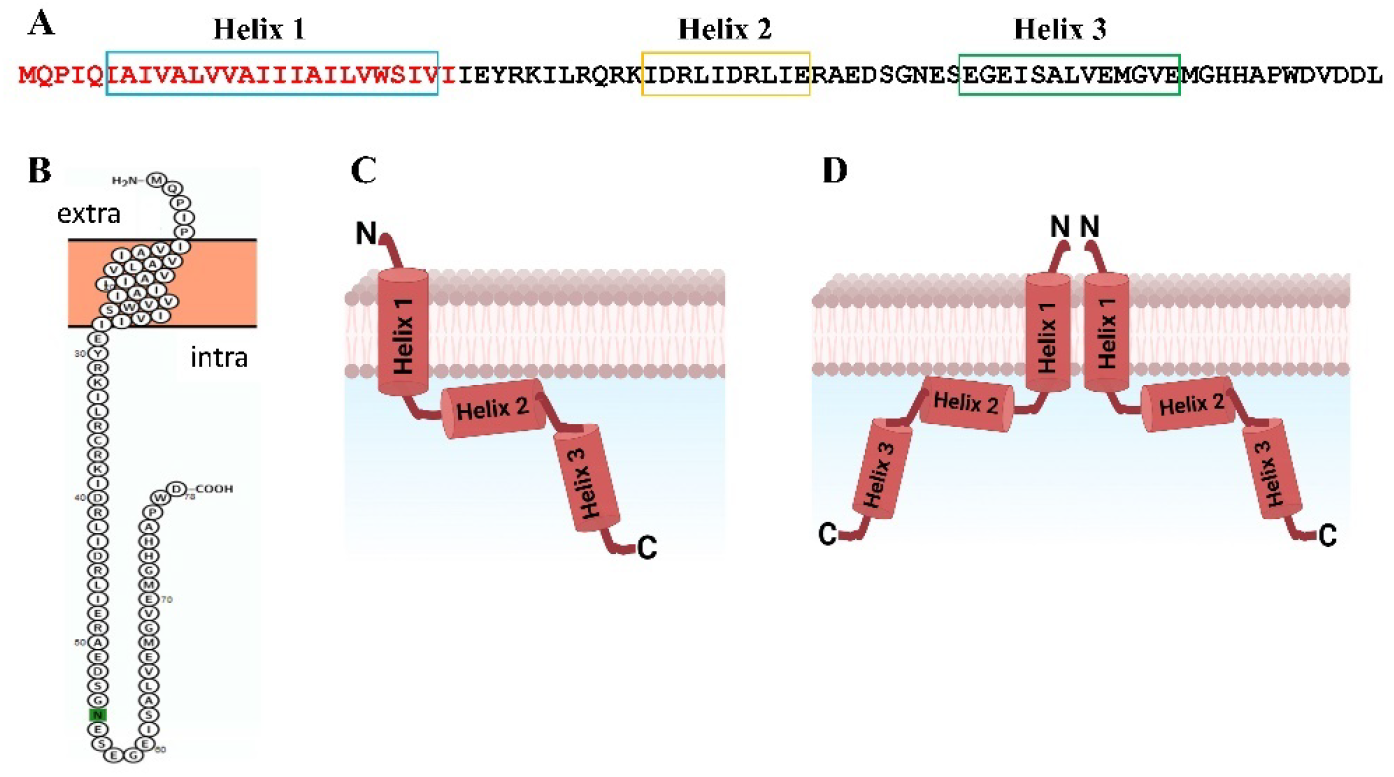
Amino acid sequence of Vpu protein and its topology in lipid membranes: (A) Amino acid sequence of HIV-1 Vpu is shown—The N-terminal is in red and the C-terminal is in black; the amino acids forming Helix 1 (TMD), Helix 2 and Helix 3 are shown in boxes. (B) Monomer organization predicted by *PROTTER* program is shown; (C) Proposed monomer organization is shown with the N-terminal Helix 1 traversing the lipid bilayer, Helix 2 interacting with membrane surface and highly soluble Helix 3; (D) Vpu oligomerization via the transmembrane Helix 1 is shown (only two monomers are shown for clarity).

Through its N-terminal Helix 1 (TMD), Vpu oligomerizes in lipid environment forming a pentameric ion-conducting channel.(9-11) Thus, its monomer structure organization and self-oligomerization classify Vpu as a member of viroporins family.(4,12) These proteins are encoded by almost all human health-threatening viruses; they support virus adaptation, proliferation and pathogenicity; and are targets for drug development.(12-15) In the infected cells, Vpu has two major biological functions, which are fulfilled by its N- and C-terminal domains: (i) modification of signaling pathways, mostly through the C-terminal interactions with host proteins resulting in downregulation of the CD4 receptor; and (ii) enhancement of newly formed virions release, mostly due to the viroporin activity through the N-terminal homo-oligomer.(16-18) These functions greatly facilitate HIV-1 infection by either cell-to-cell or cell-free spread.(19,20)

Because of diverse functions of Vpu that are critical to HIV-1 adjustment and proliferation, thorough knowledge about its structure-function relationship at the molecular level could inform on approaches to regulate the function of this protein. Yet, to date the structural information about the full-length (FL) Vpu in its monomeric and oligomeric forms is rather limited. It is also not well understood how Vpu interacts with lipid membranes to form self-oligomers (presumably pentamers), or participates in interactions via its TMD or C-terminal region with host proteins.

Here, we report our findings about the oligomerization forms of FL Vpu in solution and lipid environment. For the first time to the best of our knowledge, we observed the formation of FL Vpu pentamer and co-existence of multiple oligomeric structures as well as a monomer of Vpu in membrane environment. These studies were possible owing to our success in producing the Vpu protein in soluble form, which was aided by the design of a fusion construct containing maltose binding protein (MBP) moiety followed by a short linker and the FL Vpu (MBP-Vpu chimera protein). We produced large quantities of highly pure MBP-Vpu protein and further conducted structural studies to assess the oligomerization state of this protein. To this end, we utilized analytical size exclusion chromatography (SEC) and negative staining transmission electron microscopy (nsEM).

## RESULTS

### Design, cloning and production in soluble form of MBP-Vpu fusion protein

We engineered a construct of MBP fused to the N-terminal of FL Vpu (MBP-Vpu protein), and cloned it in a bacterial expression vector (pET15b) resulting into the MBP-Vpu/pET15b plasmid (Figure 2).

**Figure 2.**
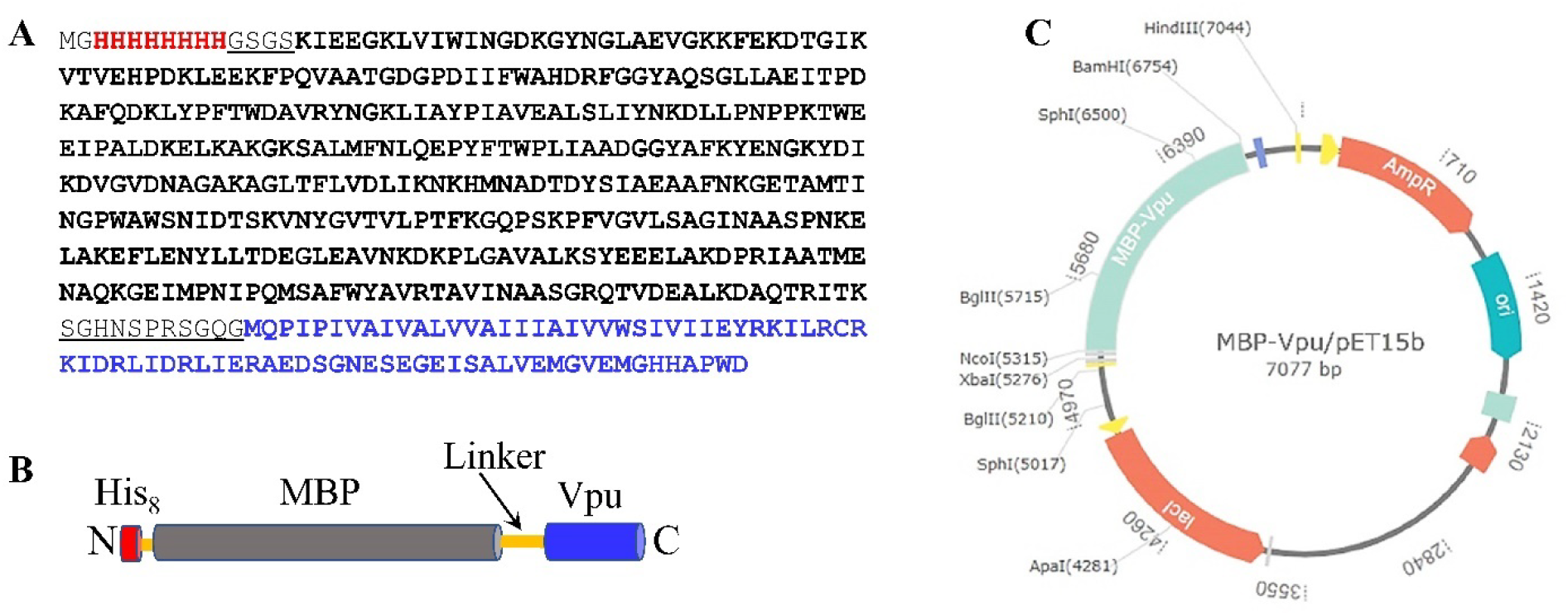
Design and cloning of MBP-Vpu protein construct: (A) The amino acid sequence of the FL MBP-Vpu fusion protein is shown—The His_8_ tag is in red, MBP is in black, Vpu is in blue, and the linkers His_8_-MBP and MBP-Vpu are underlined. (B) Schematic representation of the MBP-Vpu protein is shown—The color code is as in (A) except the linkers, which here are shown in orange. (C) MBP-Vpu plasmid map is shown—The gene encoding the MBP-Vpu protein was cloned in pET15b vector at NcoI/BamHI restriction sites.

The design of MBP-Vpu construct was carefully planned to have a His_8_ affinity purification tag, which allows for more efficient Ni^2+^ affinity purification compared to the standard His_6_ tag.(21) Further, the utilization of MBP was multi-directional—We wanted to increase the solubility of the MBP-Vpu construct, which indeed enabled us to produce large milligram quantities of the protein in soluble form in *E. coli*; MBP was also used as an affinity purification tag; and another purpose was to increase the effective molecular weight and size of Vpu fusion protein, making it suitable for visualization by EM. For the last purpose, we kept the linker between MBP and Vpu relatively short, i.e. 11 amino acids, so the final MBP-Vpu oligomer would be sufficiently conformationally stable to allow further imaging using electron microscopy (EM) and possibly solving the high-resolution structure of this oligomer. Such design to study small proteins and their complexes in lipid or just buffer environment has gained high popularity and success in the last years. (22,23)

We transformed this plasmid into *E. coli* cells and the expressed protein was indeed found in the soluble cytosolic fraction. This enabled us to handle the MBP-Vpu as a soluble protein in the following purification procedures. The double affinity tag (His_8_ plus MBP) was very beneficial for the purification process and we obtained highly pure protein by utilizing consecutively Ni^2+-^ affinity and amylose-affinity chromatography.

The calculated molecular weight and molar extinction coefficient (ε) of our MBP-Vpu protein are 52.1 kDa and 78,840 M^-1^cm^-1^, respectively (based on *ExPasy Prot Param* software). This extinction coefficient was used to estimate the protein molar concentration in the subsequent experiments.

The high protein purity was confirmed by SDS-PAGE and WB (Figure 3). Both, SDS-PAGE and WB detected multiple protein bands and the band multiplicity was reproducible in several experiments. These bands are most likely due to the formation of heterogeneous protein oligomers in SDS environment. This is quite feasible, as SDS is a membrane mimetic.(24) Thus, bands at ca. 50 kDa, ca. 100 kDa, and ca. 250 kDa were detected corresponding to MBP-Vpu monomer, dimer and pentamer, which is consistent with the oligomerization of MBP-Vpu in the membrane mimetics discussed later. The apparent bands at lower than 50 kDa molecular weight might be a result of protein degradation, but more likely they are a result of anomalous protein migration due to smaller effective size of folded in SDS FL Vpu protein. The absence of protein degradation was also confirmed by the SEC results on MBP-Vpu in buffer, also discussed below.

**FIGURE 3.**
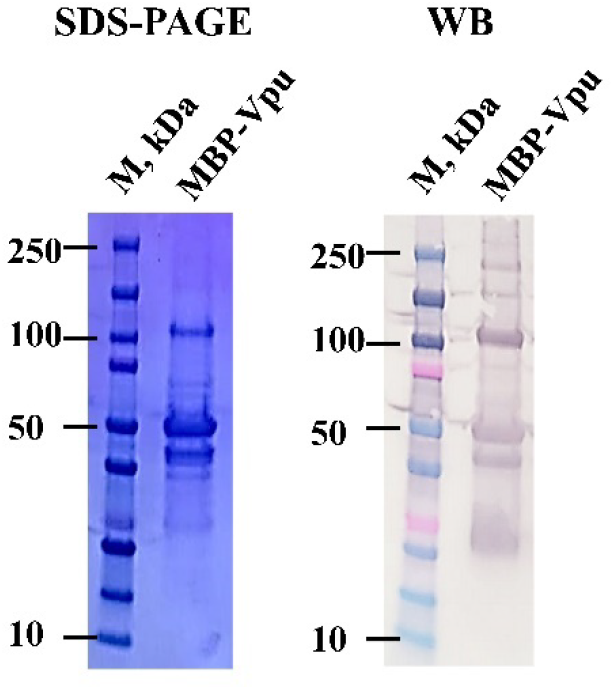
SDS-PAGE and western blotting (WB) of double affinity purified MBP-Vpu. We observed several protein bands: all bands detected by SDS-PAGE were WB positive indicating they all are due to MBP-Vpu.

An MBP-FL Vpu construct was previously created and investigated, but mostly in context of *in-cell* experiments than as a tool for *in vitro* structural studies.(5) Therefore, we believe our unique MBP-Vpu construct is the first to yield insights from *in vitro* studies on the structure and oligomer formation of HIV-1 Vpu protein. Also, MBP was utilized as a fusion tag in studies of Vpu C-terminal interactions with host proteins,(18) making it a proven tool for study of this type of viral proteins.

### MBP-Vpu forms protein concentration-dependent oligomers in aqueous solution with no lipid mimetic present

Literature strongly suggests that Vpu forms oligomers in the native membranes of Golgi, ER and plasma membrane, and in model membrane mimetics as well. (4,5,9-11) Therefore, our initial intention was to produce FL Vpu in a soluble form as a fusion construct with MBP and then reconstitute it in lipid environment for functional and structural investigations. Studying the structure of Vpu in aqueous non-membrane context was indeed not on our initial plan. Vpu has highly hydrophobic regions (TMD) and in general has not been anticipated to acquire any thought-provoking structure outside of membrane. The soluble Vpu C-terminal, was produced previously to investigate its interaction with AP-1 complex; and this study suggested that Vpu binds several subunits of the AP-1 complex as a monomer.(18) However, when we conducted SEC on our highly purified MBP-Vpu, surprisingly, we found that this protein elutes predominantly as a high-molecular weight oligomer (Figure 4A). In some cases, a second much less intensive elution peak corresponding to lower-molecular weight protein was observed as well (Supplementary Figure 1). We collected the SEC fractions corresponding to the main protein elution peak and subjected them to SDS-PAGE and WB. The results from this experiment explicitly confirmed that the elution peak is of MBP-Vpu protein (Figure 4B). Next, we run SEC using a control sample containing mixture of soluble proteins with known molecular weight (SEC protein standard) to estimate the molecular weight of the MBP-Vpu oligomers (Figure 4A). We found that the molecular weight of the oligomer, which elutes between 8.5 ml and 10.5 ml, is greater than 200 kDa, as the elution peak of this protein occurs before but close to the elution peak of β-amylase from sweet potato (200 kDa) (Figure 4A). This result unambiguously demonstrates that MBP-Vpu forms stable oligomers in solution. These MBP-Vpu oligomers are most likely pentamers with molecular weight of ca. 260 kDa, which is five times greater than the molecular weight of MBP-Vpu monomer, i.e. 52.1 kDa. Moreover, the low-intensity peak at 15-17 ml in the SEC elution profile (Supplementary Figure 1) corresponds to MBP-Vpu monomer, as it occurs between the peaks of albumin from bovine serum (66 kDa) and carbonic anhydrase from bovine erythrocytes (29 kDa) from the SEC standards; the protein in this peak is also WB-positive. Strikingly, we did not observe any intermediate oligomers of MBP-Vpu in buffer solution, suggesting a monomer-to-pentamer equilibrium under the conditions of this experiment, i.e. 50 µM protein monomer in buffer with 10% glycerol.

**Figure 4.**
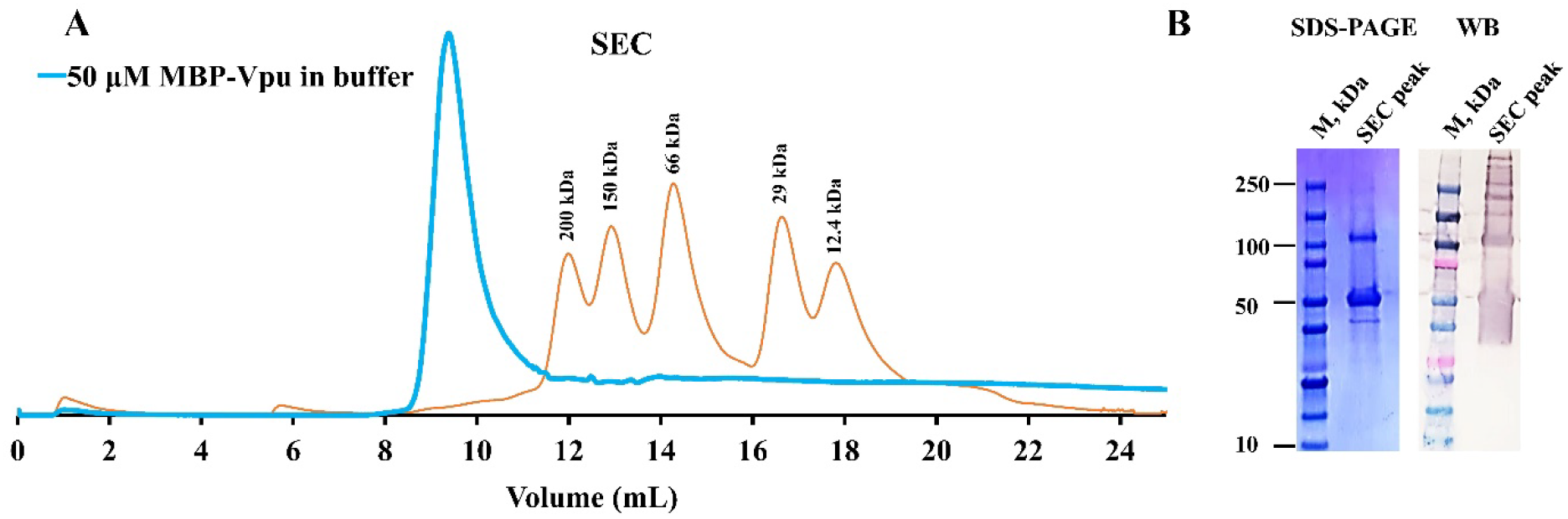
SEC data confirm that MBP-Vpu forms pentamers in solution: (A) The SEC data for MBP-Vpu are shown in blue. An elution peak at 8.5 ml to 10.5 ml was observed. The fractions corresponding to this peak were pooled and subjected to SDS-PAGE and WB. The SEC data for mixture of protein standards is shown in red with the protein molecular weight indicated next to each elution peak (β-amylase from sweet potato – 200 kDa, alcohol dehydrogenase from yeast – 150 kDa, albumin from bovine serum – 66 kDa, carbonic anhydrase from bovine erythrocytes – 29 kDa, and cytochrome c from horse heart – 12.4 kDa). (B) SDS-PAGE and WB results for the pooled MBP-Vpu fractions as described in (A) are shown—The protein in this elution peak was WB positive, confirming that it is MBP-Vpu.

We wanted to find out which region of MBP-Vpu is responsible for the oligomerization in solution. MBP does not self-oligomerize in solution and when is used as a fusion tag.(25,26) On the other hand, the TMD region of Vpu is responsible for oligomerization in lipid environment, (9,11). So, it may well be that this region contributes to the observed protein self-association in aqueous solution possibly due to the hydrophobic effect to minimize the contact with the water solvent.(27) We further examined the Vpu C-terminal (residues 28-81 in FL Vpu), and found that in solution it is predominantly a monomer (Supplementary Figure 2), although it was earlier suggested that this region is important for oligomer stabilization in lipid;(10) in addition, self-association in buffer solution was previously observed for the cyto domain of gB fusogen from herpes simplex virus,(28) indicating that juxtamembrane soluble regions of single-pass transmembrane proteins might pay stabilizing role in their oligomerization.

We next utilized nsEM to visualize these MBP-Vpu oligomers in solution. Certainly, our goal to increase the effective molecular weight and size of Vpu by fusing it to MBP was very fruitful and we successfully observed well-defined oligomeric structures at 5 µM MBP-Vpu (Figure 5A) and 10 µM MBP-Vpu (Supporting Figure 3). These protein assemblies were similar in size, and were about 20 nm in diameter when observed from the top of the oligomer (Figure 5A). The uniformity of protein aggregates was also confirmed by the particle size analysis, which we conducted—Among a set of more than 30 protein particles, we estimated high particle homogeneity with average diameter of ca. 19.8 nm (Supporting Figure 4).

**Figure 5.**
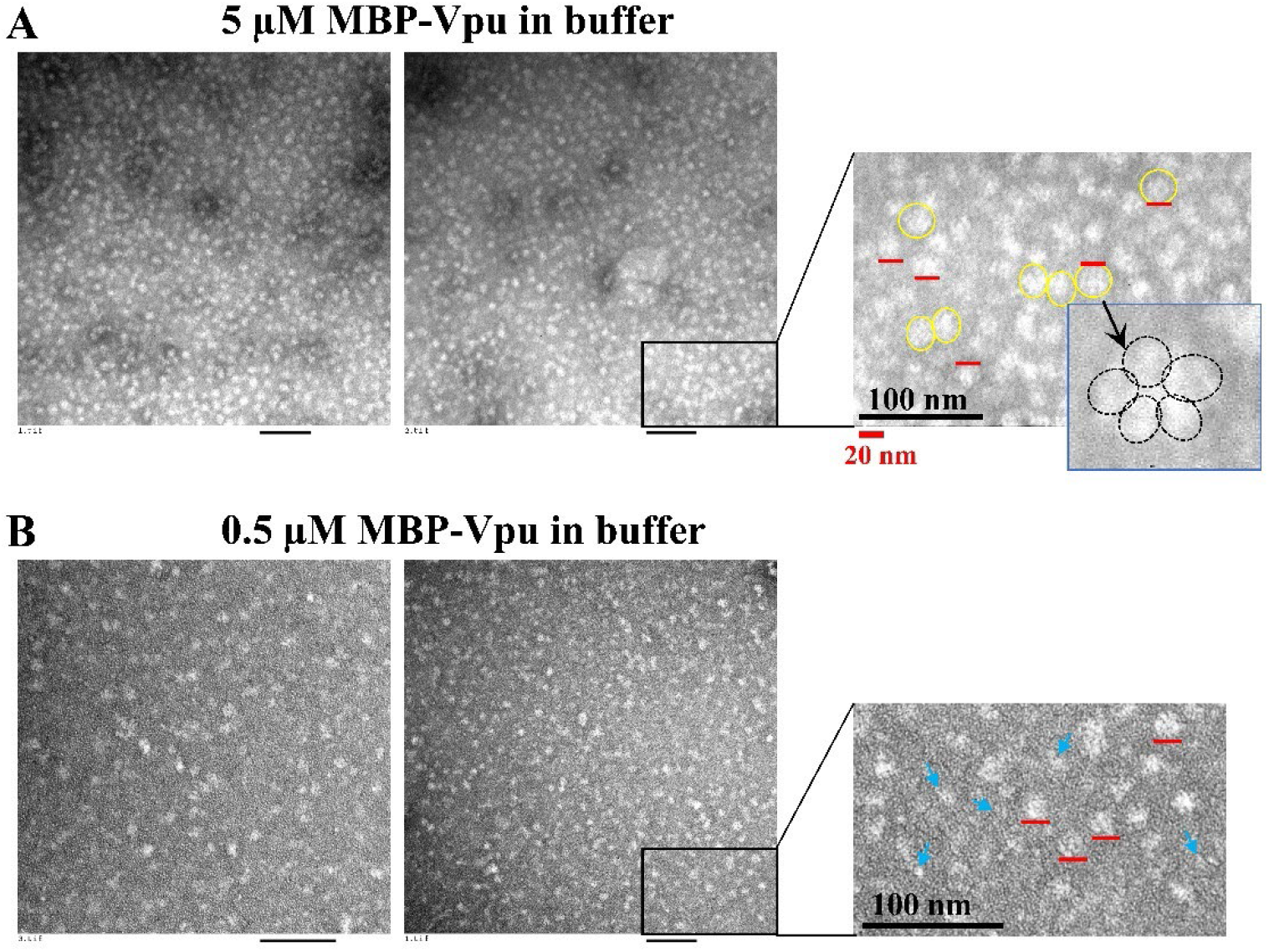
Representative EM images of negatively stained MBP-Vpu in solution: (A) Data for 5 µM protein are shown. (B) Data for 0.5 µM protein are shown. In all images, the black bars correspond to 100 nm and red bars correspond to 20 nm. It is apparent from the figure that at the higher protein concentration of 5 µM more homogeneous oligomers are formed with the diameter of their top side of ca. 20 nm (shown in yellow circles in (A)). The coarse estimate is that these oligomers are pentamers (inset in the rightmost panel in (A)). The blue arrows in (B) show smaller protein aggregates, which are present in the sample of 0.5 µM MBP-Vpu.

Coarse analysis shows that the pentamer is the most probable oligomeric state in the observed MBP-Vpu aggregates (Figure 5), which is in agreement with SEC data and literature for Vpu in lipid.(9,11) This is further reinforced by the consideration that the empty of substrate MBP monomer is about 6.8 nm long, based on its crystal structure,(29) so if added together the possible size of Vpu pentameric pore, the length of MBP-Vpu linker and the size of MBP monomer, observing pentameric structures with 20 nm diameter is quite plausible (Supporting Figure 5). We additionally observed that the oligomer stability was affected by protein dilution, since when the protein concentration was 0.5 µM, oligomers with reduced size than the those formed at 5 µM protein concentration were observed (Figure 5B); MBP-Vpu oligomers were possibly also present. This indicates that the MBP-Vpu oligomer assembly is protein (monomer) concentration dependent. However, this concentration dependence is valid only for very low micromolar to nanomolar protein concentration. At higher protein concentration, fully assembled and stable oligomers are formed, i.e. the process is possibly cooperative.

Altogether, our results from SEC and nsEM strongly suggest that MBP-Vpu protein forms oligomers in solution most likely via Vpu TMD.

### In lipid environment, MBP-Vpu oligomerization and oligomer morphology depend on the nature of lipid membrane mimetic and protein-to-lipid (detergent) molar ratio

After revealing that MBP-Vpu forms stable oligomers in solution, we were interested to know how the oligomeric state of this protein is affected by the addition of lipid membrane mimetic. We reconstituted the soluble MBP-Vpu in three different lipid-like environments that are β-DDM (n-Dodecyl-beta-Maltoside), lyso PC/PG (14:0 lyso PC/14:0 lyso PG) mixture at 50:50 mol% and DHPC/DHPG (1,2-dihexanoyl-sn-glycero-3-phosphocholine/1,2-dihexanoyl-sn-glycero-3-phospho-(1’-rac-glycerol) (sodium salt) mixture at 50:50 mol%. All three lipid mimetics form micelles of different sizes and shapes.(30-32) However, we used DHPC/DHPG mixture at 2 mM or less, which is below the lipid critical micelles concentration (CMC) of about 15 mM, since concentrations at and above CMC produced difficult to interpret nsEM data. The lipid membrane mimetics were selected due to their wide use in membrane protein studies,(32-37) and lipids with negatively charged headgroups (lyso PG and DHPG) were included, since they are deemed important for interactions with positively charged residue in viroporins.(12)

We conducted analytical SEC experiments on MBP-Vpu in buffer supplemented with 1 mM β-DDM or 1–2 mM lyso PC/PG. In both cases the protein concentration used was 25–50 µM. Strikingly, in both detergent/lipid environments, we detected several SEC elution peaks. In β-DDM we observed elution peaks at 8.5–10.5 ml, as well as with peak maxima at 12 ml and close to 16 ml (Figure 6A), with the peak at 12 ml being the most intensive under our experimental conditions of 50 µM protein. Similar SEC profile occurred for protein in lyso PC/PG, but the intensities of the elution peaks for different protein oligomeric states depended also on the protein concentration and additionally shoulders at 7–8.5 ml and 12.5–14.5 ml appeared but the peak at 16 ml was not present (Figure 6B). In comparison to the elution profile of protein standards with known molecular weight (Figure 4A), the coarse estimate is that the peak at 8.5–10.5 ml corresponds to protein with molecular weight of 260 kDa, representing MBP-Vpu pentamers, the peaks from 10.5 ml to 12.5 ml correspond to protein with molecular weight about 200 kDa, suggesting MBP-Vpu tetramers, and the last peak at 16 ml is of MBP-Vpu monomer. Thus, our results point to significant restructuring of MBP-Vpu protein in lipid environment. However, this restructuring is not in the direction of stabilizing a unique protein oligomer. Instead, greater oligomer polymorphism occurred when Vpu transitioned from aqueous to its native membrane-like environment.

**Figure 6.**
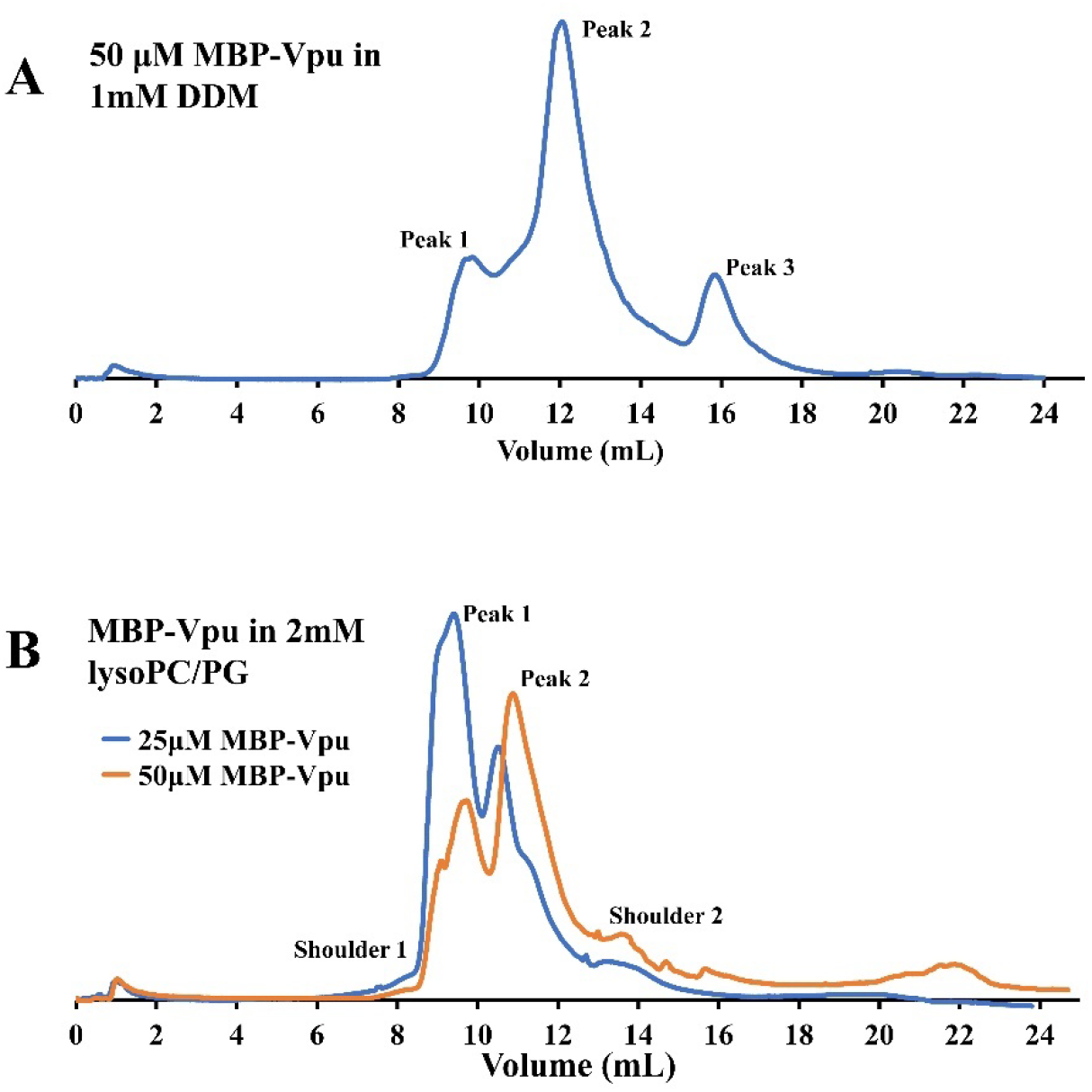
SEC data for MBP-Vpu in β-DDM and lyso PC/PG: Data for 50 µM protein in 1 mM β-DDM are shown; (B) Data for 50 µM and 25 µM protein in 2 mM lyso PC/PG are shown. In both environments, multiple elution peaks of MBP-Vpu corresponding to distinct protein oligomeric states were detected.

We performed nsEM to visualize the MBP-Vpu oligomers in β-DDM, lyso PC/PG and DHPC/DHPG. We used a range of protein and detergent/lipid concentrations that were: 5 µM and 1 µM protein both in 1 mM *β*-DDM (protein-to-detergent molar ratios, P/Ds, of 1:200 and 1:1000, respectively) (Figure 7); 5 µM protein in 1 mM lyso PC/PG, 5 µM protein in 2 mM lyso PC/PG, 0.5 µM protein in 2 mM lyso PC/PG (protein-to-lipid molar ratios, P/Ls, of 1:200, 1:400 and 1:4000, respectively) (Figure 8); and 1 µM protein in 1 mM DHPC/DHPG, 1 µM protein in 2 mM DHPC/DHPG, and 5 µM Protein in 2 mM DHPC/DHPG (P/Ls of 1:1000, 1:2000, and 1:400, respectively) (Figure 9).

**Figure 7.**
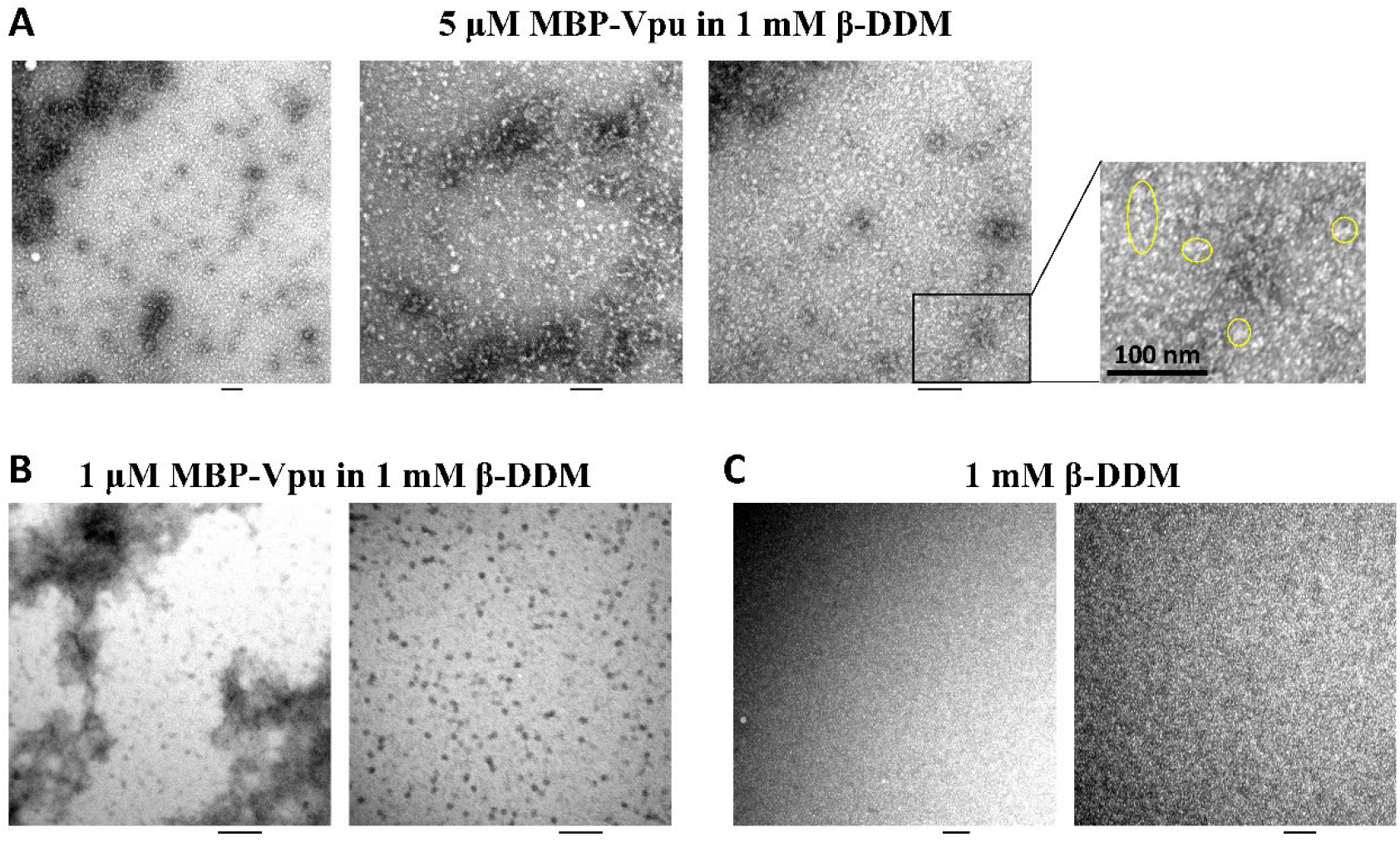
Representative nsEM images of MBP-Vpu in β-DDM: (A) data for 5 µM protein in 1 mM β-DDM are shown; (B) data for 1 µM protein in 1 mM β-DDM are shown; and (C) data for control sample of 1 mM β-DDM are shown. Heterogeneous aggregates, including linear arrays of MBP-Vpu, are encircled in yellow. The 100 nm bar is shown in black.

**Figure 8.**
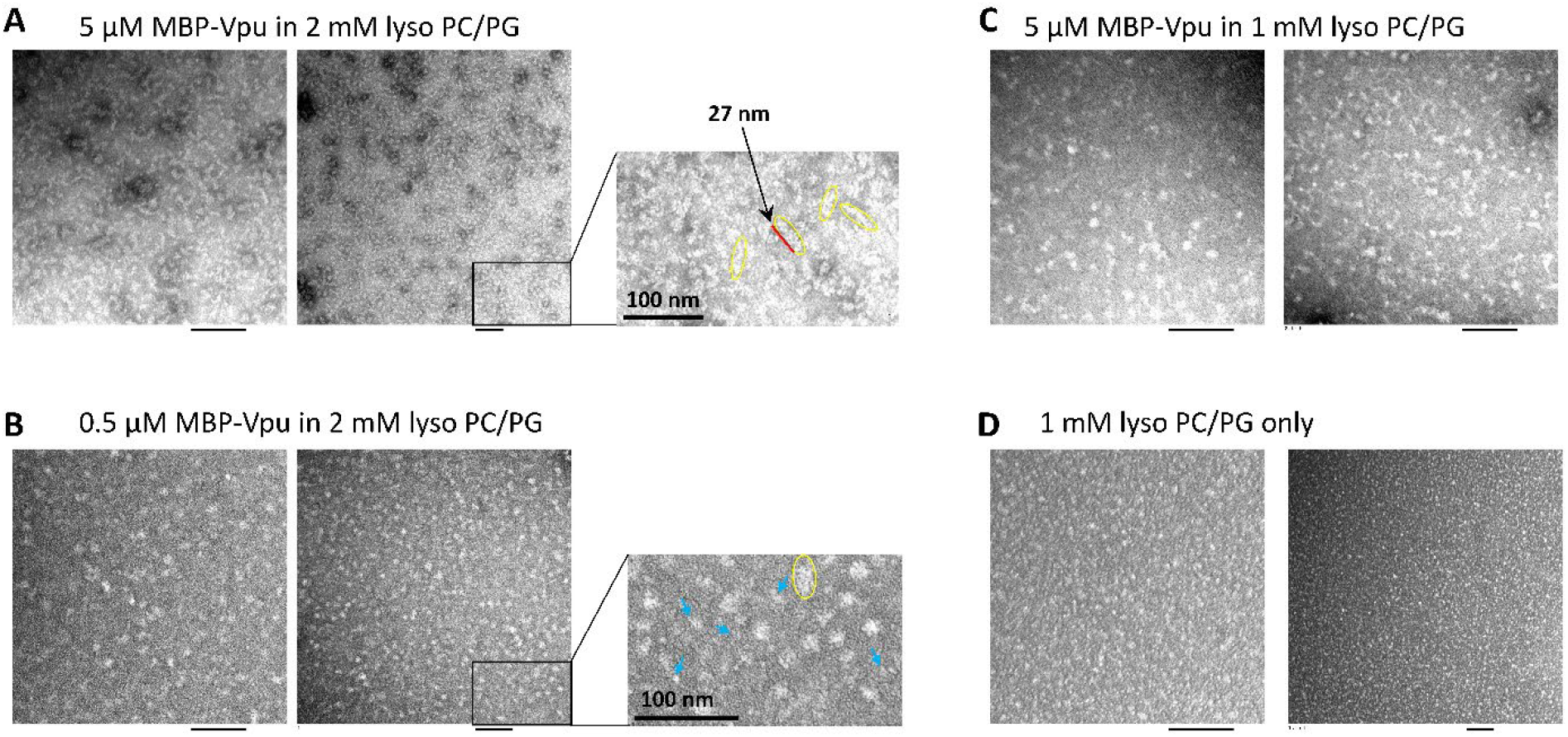
Representative nsEM images of MBP-Vpu in lyso PC/PG: (A) data for 5 µM protein in 2 mM lyso PC/PG are shown; (B) data for 0.5 µM protein in 2 mM lyso PC/PG are shown; and (C) data for 5 µM protein in 1 mM lyso PC/PG are shown; (D) data for control sample of 1 mM lyso PC/PG are shown. The 100 nm bar is shown in black. The formation of linear MBP-Vpu oligomers is clearly visible in (A), whereas greater protein aggregates were observed in (B) and (C). The linear MBP-Vpu aggregates are enclosed in yellow and the smaller aggregates are indicated with blue arrows. The red bar in (A) rightmost panel shows 27 nm linear protein aggregate.

**Figure 9.**
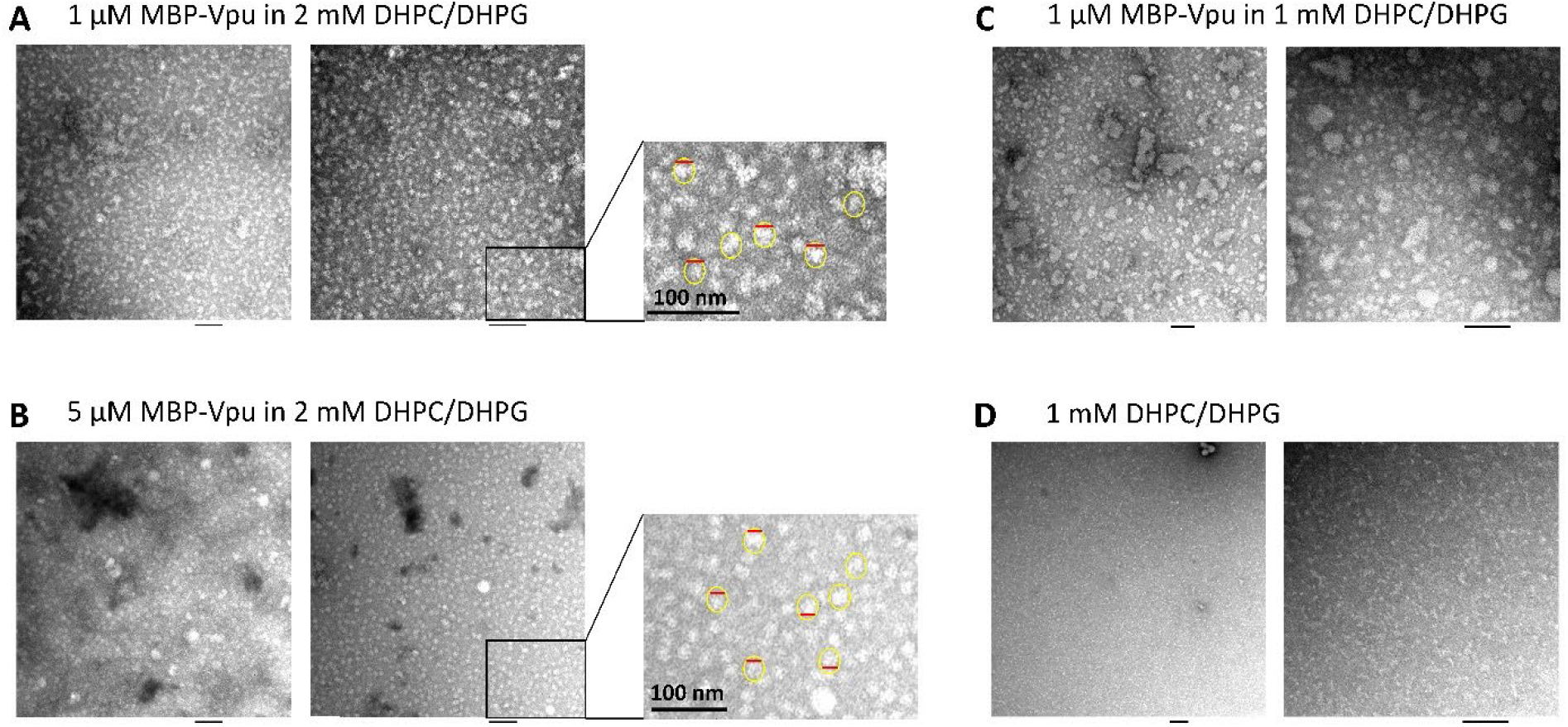
Representative nsEM images of MBP-Vpu in DHPCDH/PG: (A) Data for 1 µM protein in 2 mM lipids are shown; (B) Data for 5 µM protein in 2 mM lipids are shown; (C) Data for 1 µM protein in 1 mM lipids are shown; (D) Data for control sample of 1 mM lipid are shown. The aggregates that are similar to MBP-Vpu pentamers in just buffer are encircled in yellow and the 20 nm bar is in red. The 100 nm bar is shown in black.

In β-DDM, we found that MBP-Vpu oligomeric state is highly heterogeneous with aggregation order increasing upon increasing the protein concentration, i.e decreasing P/D (Figure 7A vs. 7B). Remarkably, this indicates that the Vpu oligomers, more specifically the pentamers, possess reduced stability in lipid-like environment compared to the oligomers in aqueous environment, i.e. the self-attraction of Vpu monomers in lipid environment is weaker. Thus, in the case of 1 µM protein (P/D of 1:1000) we observed mostly dispersed low-order oligomers and possibly monomers (which are too small in size to be clearly distinguished by the technique used). At 5 µM MBP-Vpu (P/D of 1:200) larger protein oligomers persisted (encircled in yellow in Figure 7A), but they were apparently smaller than pentamers. What is more, under the P/D of 1:200 condition, elongated linear-like aggregates of MBP-Vpu were also formed (surrounded by yellow ellipse in Figure 7A), representing novel structural organization of membrane-bound Vpu, which has not been reported previously.

MBP-Vpu oligomers were detected in lyso PC/PG as well, but varying patterns of oligomerization were observed for P/Ls of 1:400 and 1:200 vs. 1:4000. Remarkably, for P/L of 1:400, similarly to P/D 1:200, we observed the formation of close to liner MBP-Vpu arrays. These linear protein aggregates were well-pronounced in lyso PC/PG environment with a length of up to or even longer than 30 nm (Figure 8A) than that in β-DDM (Figure 7A). To confirm this lipid-driven protein organization, we further conducted EM on negatively stained MBP-Vpu linked to 5 nm gold nanoparticles (GNPs) in lyso PC/PG (we reproduced the 5 µM protein in 2 mM lyso PC/PG conditions). Of note is that the concentration of GNPs in this sample was 0.04 µM (due to very low concentration of GNPs stock solution), which is significantly lower than the protein concentration, so not every MBP-Vpu monomer was linked to a GNP. Nevertheless, we could again clearly trace the GNP-MBP-Vpu arrays (Supporting Figure 6). Thus, in lipid-like environment, we observed for the first time an unexpected, still, most likely physiologically relevant organization of Vpu protein. Once more, despite the different protein concentration ranges, the data from nsEM agree well with the SEC results showing multiple and broad elution peaks (Figure 6B). More heterogeneous MBP-Vpu aggregates were detected for P/Ls of 1:4000 and 1:200 (Figure 8B,C). Still, along with shorter and more round-shaped protein oligomers, we also observed elongated aggregates as for P/L 1:400.

Large variations in MBP-Vpu oligomers were detected by nsEM in DHPC/DHPG as well (Figure 9). The most homogeneous and well-shaped oligomers were observed for samples of 1 µM or 5 µM protein in 2 mM lipid (P/L of 1:2000 and 1:400) (Figure 9A,B), validating further the finding that the protein concentration plays a role in Vpu oligomer assembly in lipid environment.

All in all, the results from experiments using both SEC and nsEM techniques strongly suggest that membrane environment modulates the strength of Vpu-Vpu monomers vs. Vpu-lipid attraction. Furthermore, it seems that the type of lipid mimetic used, i.e. mild non-charged detergent (β-DDM) vs. mixture of non-charged/charged lyso lipids (lyso PC/PG) vs. mixture of true non-charged/charged short-chain lipids (DHPC/DHPG), has an important role in defining the quaternary structure of Vpu, but protein concentration also plays a role. It is worth mentioning that the Vpu oligomer assembly was more efficient in closer to native lipid environment (lyso PC/PG and DHPC/DHPG) compared to detergent (β-DDM), possibly reflecting this protein properties in native membranes. Previously, such an effect of detergent vs. lipid on influenza M2 protein self-associating was also identified.(34)

## DISCUSSION

Vpu plays an important role in HIV-1 life cycle through its function as an ion-conducting channel in host membranes owing to the protein TMD oligomerization (Figure 1) and interaction of its soluble C-terminal domain, as well as TMD, with host proteins.(4,5,9-11) In addition to its involvement into transporting ions across cellular membranes, viral egress and budding,(38,39) etc., Vpu was found to suppress cellular signals critical for proper immune response in already infected cells.(40) In spite of the extensive studies to elucidate the structure-function relationship of this protein, the success thus far has been moderate. NMR spectroscopy and X-ray crystallography provided insights into the structure of isolated C-terminal region (11,18) and TMD.(11) However, currently, there is no clear understanding of what is the structural basis of Vpu association with host proteins and details about how this protein interacts with lipid environment were not known. No any structure of FL Vpu has been reported to date. Even though the prevalent opinion that Vpu organizes and functions as a pentamer, (11) up to now, no data showing the fully assembled pentamer of FL Vpu has been provided. Also, it was proposed that Vpu forms pentamer in membranes of Golgi and intracellular vesicles, but not in the endoplasmic reticulum (ER),(5) suggesting that functional Vpu exists in multiple quaternary structures, possibly each of these structures corresponds to a distinct protein function. Still, these multiple structures and their involvement in viral processes in the host cell are yet to be fully characterized. This lack of comprehensive knowledge about molecular properties of Vpu presents a serious obstacle in acquiring complete understanding about this protein multiple roles in HIV-1 adaptation and persistence and in the development of anti-Vpu pharmaceuticals.

To contribute to the overcoming of this deficiency, we undertook a study to assess the Vpu oligomerization in lipid environment by using a combination of analytical SEC and EM. Due to the small size of FL Vpu, ca. 9 kDa, conducting EM on this protein even in its pentameric form would be difficult due to the current limitations for high-resolution structure determination of membrane proteins by EM. Therefore, we selected to benefit from the lately developed fusion construct methodology to visualize small proteins in membrane by generating a protein chimera linking a larger and stable soluble protein to one of the termini of the small membrane proteins and thus increasing the effective protein size/molecular weight.(22) To do so, we generated a fusion construct of MBP linked to the N-terminal of FL Vpu (Figure 2A,B). The linker of 11 amino acids between the two proteins was carefully selected to minimize the protein flexibility and at the same time to avoid protein oligomerization hindrance due to presence of bulkier MBP.

Initially, we subjected the highly-pure MBP-Vpu protein in buffer, with no lipid or detergent added, to analytical SEC and nsEM and surprisingly discovered that the protein indeed forms stable pentamers (Figures 4 and 5) in just aqueous environment. Both SEC and nsEM confirmed that above 5 µM concentration (used in EM experiments) and up to 50 µM concentration (used in SEC experiment), probably even higher concentrations, MBP-Vpu existed exclusively in pentameric form. Even more remarkable is that within this concentration range a single transition from monomer (low abundance) to pentamer (high abundance) was observed (Supplementary Figure 1) suggesting a monomer-to-pentamer transition. Based on prior studies,(25,26,41) it is well known that MBP itself does not form oligomers in solution. Therefore, we believe the observed oligomerization is via Vpu moiety of the protein, namely its TMD.

The physiological relevance of Vpu pentamer in solution is currently unclear, as Vpu has been known only as a single-pass transmembrane protein. It may well be that the assembly of such an oligomer is just a result of hydrophobic effect driving the TMDs together to minimize the contact with hydrophilic environment; the stability of the whole complex could be because of the large MBP portion contributing to the significant hydrophilic nature to the MBP-Vpu fusion protein. However, even more fascinating is the pentameric organization of MBP-Vpu in solution with well-ordered in a shape close to filled circle MBP moieties when looking from the top of the oligomer (Figure 5 and Supplementary Figure 4). This protein behavior likely suggests that the pentameric form of Vpu is optimal for oligomer stability and is amino acid sequence-defined. Strikingly, the oligomeric order in solution corresponds to that proposed for Vpu channel/pore in lipid membranes. Thus, by the mean of analogy, the pentamer in solution could be considered as a model of Vpu channel in lipid environment. Still, one could argue that the forces stabilizing the structure of Vpu pentamer in solution and in lipid environment are quite different, therefore distinct folds of FL protein monomer within the pentamers in these two environments cannot be ruled out. More studies to possibly determine the high-resolution structure of Vpu pentamer would be beneficial to better understand the observed result.

We further studied the oligomerization profile of MBP-Vpu reconstituted in lipid membrane mimetics that are β-DDM, lyso PC/PG and DHPC/DHPG. Large protein oligomer polymorphism was detected by both SEC and nsEM in all three membrane environments (Figures 6, 7, 8 and 9). What is more, it seems that the transition from hydrophilic to membrane environments led to substantial protein restructuring including destabilization of the stable pentamer in solution. Indeed, the oligomeric helical proteins in lipid bilayers participate in complex and even competing contacts including helix-helix and helix (amino acid)-lipid interactions.(42) Thus, it seems that the Vpu monomer-monomer stabilizing contacts outside of the native aquaphobic environment are now replaced to a certain extent by hydrophobic protein-lipid interactions, which lifts the mandate for pentameric organization. Still, the MBP-Vpu pentamer was observed in membrane environment, pointing that this is definitely one of the functional native membrane-bound Vpu forms, possibly representing the Vpu viroporin (ion-conducting channel/pore). Furthermore, it is likely that the protein monomeric state is physiologically relevant as well, as previous studies suggest that Vpu does not form oligomers in ER,(5) and Vpu interacts with clathrin adaptor protein complex 1 as a monomer, (18) although in the latter study a truncated version of the Vpu C-terminal domain was used.

The most striking result about MBP-Vpu in lipid membranes is the observation of MBP-Vpu liner aggregates (arrays) when the protein was reconstituted in lyso PC/PG at P/L 1/400 (Figure 8). Such Vpu structural organization has not been reported before and linking it to a specific function is currently quite imprecise. It is plausible that this Vpu organization affects membrane morphology via destabilizing effect.

In conclusion—We generated an MBP-Vpu construct, which was beneficial to produce the protein in soluble form, as well as to observe protein oligomerization pattern in both aqueous and membrane environment. We propose that the observed in just buffer MBP-Vpu oligomer could be subjected further to high-resolution structural determination and used as a model of Vpu channel/pore in lipid membranes. We also detected multiple oligomeric forms of MBP-Vpu in lipid environments, each of which most likely represents a functional state of Vpu, depending on the particular role the protein fulfills. Figure 10 summarizes our view of how Vpu supramolecular structure relates to its function. We believe our findings will stimulate further studies to comprehensively characterize Vpu structures leading to better understanding of this protein function.

**Figure 10.**
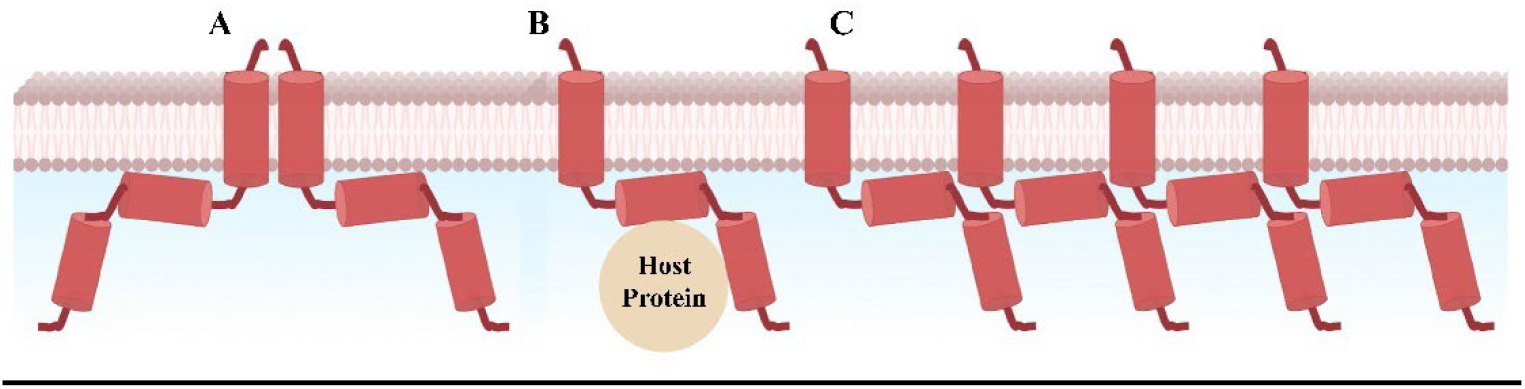
Proposed Vpu organization in cellular membranes: (A) Vpu forms pentameric structures possibly with ionic channel/pore activity (only two subunits are shown for clarity); (B) Vpu exists and functions as a monomer (likely participates in interactions with host proteins via both its TMD and C-terminal domain); (C) Vpu forms linear aggregates with currently not well understood function.

## EXPERIMENTAL PROCEDURES

### Design and cloning of MBP-Vpu fusion construct

We utilized protein engineering to generate a fusion construct of maltose binding protein (MBP) followed by a short linker and full-length (FL) HIV-1 Vpu for optimized structural studies, resulting into MBP-Vpu fusion protein. In addition, this construct had Hisx8 (His_8_) affinity tag at its N-terminal to facilitate protein purification. Further, we introduced a single cysteine residue at position Q36C (Figure 1), which was not utilized in the current study but will be useful in future cysteine-specific labeling and spectroscopic studies, possibly cross-linking as well. The DNA encoding this fusion protein was commercially synthesized (GenScript Inc) and cloned into pET15b bacterial expression vector at Nco I/BamH I cloning sites. The obtained MBP-Vpu/pET15b vector was further used for protein expression. We utilized the *ExPasy ProtParam* software (43) to characterize the MBP-Vpu protein properties, e.g. molecular weight, extinction coefficient, etc., and the *PROTTER version 1*.*0* program (44) to predict the transmembrane topology of MBP-Vpu.

### Expression and purification of MBP-Vpu construct

First, pET15b/MBP-Vpu plasmid was transfected into *E*.*coli* BL21(DE3) competent cells (*Lucigen*) following the recommended by the manufacturer protocol. Then, the transfected cells were plated on LB/agar/ampicillin (Amp) plate (100 µg/ml Amp) and incubated overnight at 37 °C. Then, a single colony was selected and inoculated in a flask containing 200 mL LB broth Lennox (*Sigma-Aldrich*) supplemented with 100 μg/mL Amp (*Gold Biotechnology*) and bacterial stock solution was grown in an incubator-shaker (*Innova 43/43R*) overnight for 17-18 h at 37°C. On the next day, 30 ml of the overnight cell culture was inoculated in a 3L Fernbach flask containing 2 L LB broth and 100 μg/mL Amp, and grown in an incubator-shaker at 37°C at 200 RPM for 2 h 30 min–3 h, until the absorbance at 600 nm (OD_600_) of the inoculated LB medium reached 0.70–0.80. Afterward, the temperature in the incubator-shaker was reduced from 37°C to 18°C, and IPTG (Isopropyl-β-D-1-thiogalactopyranoside) (*Gold Biotechnology*) was added to the bacterial solution to final concentration of 1mM to induce the protein expression under the control of T7 promoter. The protein expression proceeded overnight at 18°C. On the next day, cells were harvested by spinning them down in an Avanti J-15R centrifuge (*Beckman coulter;* JA-4.750 rotor) at 4,100 RPM (1,880×g) for 15 min at 4°C. The supernatant was discarded, and cell pellets were collected and resuspended in the resuspension buffer containing 20 mM HEPES (4-(2-hydroxyethyl)-1-piperazineethanesulfonic acid) (*Sigma*) pH 7.4 and 200 mM NaCl (*Sigma*)]. After that, TCEP (tris (2-carboxyethyl) phosphine) (*Gold Biotechnology*), chicken egg lysozyme (*Roche*), and PMSF (phenylmethylsulphonyl fluoride) (*Gold Biotechnology*) were added to the resuspended cell solution to final concentrations of 200 μM, 0.5-0.6 mg/mL and 1mM respectively. This solution was then subjected to sonication with a sonicator (*U*.*S. solid ultrasonic processor*) to break the cells open, and the cell debris was separated with centrifugation at 7,000 RPM (5,380 ×g) for 15 min in the same centrifuge (JA-10.100 rotor) at 4°C. The supernatant containing the soluble fraction of cell lysate was collected and subjected to double-affinity chromatography (45) to purify MBP-Vpu protein.

Double-affinity chromatography, *i*.*e*. Nickel _(_Ni^2+^, Ni)-affinity chromatography followed by amylose affinity chromatography was utilized to obtain highly pure MBP-Vpu protein. For Ni-affinity purification, Ni-NTA agarose resin (*Qiagen*) (1.5 mL/1 L cell culture) was incubated under constant agitation with soluble cell lysate in the resuspension buffer at 4°C for a duration of 1-1.5 h. Afterward, Ni-NTA agarose resin bound with protein was transferred to a gravity column and flow-through containing the unbound material was discarded. Then, to elute most of the weakly bound protein impurities, the column was washed with 10 resin volumes of buffer A containing 50 mM sodium phosphate buffer pH: 7.4, 150 mM NaCl, and 5 %(w/v) glycerol supplemented with 40 mM imidazole (Im) (*Sigma*). Subsequently, 2 resin volumes of buffer A supplemented with 300 mM Im was added to the column for the elution of the target protein, *i*.*e*. MBP-Vpu protein. Later, the protein was concentrated at 3,900 RPM (1,700 ×g) in 10 kDa MWCO ultra centrifugal filters (*Amicon*^®^) at 4°C. Protein concentration was measured using nanodrop spectrophotometer (*Thermo Scientific*) and then subjected to amylose purification technique, after removing Im by washing with an exchange Buffer B made of 20 mM Tris pH: 7.4, 200 mM NaCl, 1 mM EDTA and 5% glycerol. For the amylose affinity chromatography, pre-equillibrated with buffer B amylose resin was incubated with Ni-affinity purified MBP-Vpu protein, at 4°C for 1.5-2 h. Afterward, the amylose resin bound with protein was transferred to a gravity column and the flow-through containing the unbound material was discarded. Then, buffer B containing 20 mM Tris pH: 7.4, 200 mM NaCl, 1 mM EDTA and 5% glycerol was added to the gravity column to wash out the unbound impurities. Next, buffer B supplemented with 25 mM maltose (*Sigma*) was introduced into the column for the elution of the target MBP-Vpu protein. The flow-through was collected and the protein was concentrated at 3,900 RPM (1,700 ×g) in 10 kDa MWCO ultra centrifugal filters (*Amicon*^®^) at 4°C. Protein concentration was measured with the nanodrop spectrophotometer using the calculated extinction coefficient of 78,840 M^-1^cm^-1^. This highly-pure protein was used in further experiments.

### Sodium dodecyl sulfate gel electrophoresis (SDS-PAGE) and western blotting (WB)

The protein purity was assessed by using SDS-PAGE and WB. All the samples for SDS-PAGE and WB constituted of MBP-Vpu protein, dithiothreitol (DTT; *Sigma*), and a loading buffer (*BioRad*). The samples and marker (precision plus protein dual-color standard; *BioRad*) were loaded into 4–20% Criterion™ TGX™ precast gels (*BioRad*) immersed in Tris/glycine/SDS buffer (*BioRad*), and then electrophoresis was conducted in a midi Criterion™ vertical electrophoresis cell (*BioRad*) at 170 V. For blotting, the protein was transferred from the gel onto a 0.2 μm nitrocellulose membrane (*BioRad*) using a Criterion™ blotter (*BioRad*) overnight at 10 V and 4 °C. Protein bands on the gel were visualized by Coomassie Blue dye staining and washing with de-stain solution. We utilized colorimetric detection to visualize the MBP-Vpu band(s) on the membrane after WB transfer: Primary mouse anti-histidine tag antibody (*BioRad*) and secondary goat anti-mouse IgG antibody conjugated to Alkaline Phosphatase (*BioRad*) were used.

### Preparation of lipid and detergent stock solutions

Stock solutions of lipids were prepared prior to reconstituting MBP-Vpu in them. Lyso lipids, 14:0 lyso PC ((1-myristoyl-2-hydroxy-sn-glycero-3-phosphocholine)) and 14:0 lyso PG (1-myristoyl-2-hydroxy-sn-glycero-3-phospho-(1’-rac-glycerol) (sodium salt)) at 1:1 molar ratio were mixed in chloroform, methanol and water, the organic solvent was evaporated under the stream of N_2_ gas until no liquid was visible and then the vial was exposed to N_2_ stream for 2 more hours to remove the residual organic solvent. Thereafter, the lipid film was rehydrated for 1-2 h at 4°C in ultrapure water to obtain 50 mM total lyso lipid stock solution. In a similar manner, the DHPC (6:0 PC) and DHPG (6:0 PG) lipids at 1:1 molar ratio were mixed and a final stock solution of 40 mM total lipid in ultrapure water was prepared. These stock solutions were further used to reconstitute MBP-Vpu protein at the desired protein-to-lipid molar ratios.

All lipids were purchased from Avanti^®^ Polar Lipids, Inc.

### Assessing the MBP-Vpu oligomerization using analytical size exclusion chromatography (SEC)

We used SEC to assess the oligomerization state of the highly-pure MBP-Vpu protein. Since no other protein impurities were present, the SEC elution profile directly reports on the MBP-Vpu oligomerization state through the position of the protein elution peaks. For these experiments, we used a Superdex™ 200 increase 10/300 GL size-exclusion column (*GE Healthcare*) plugged into AKTÄ explorer 100 (Amersham Biosciences) protein purifier system. For all SEC experiments the protein buffer was exchanged to 50 mM NaPi (sodium phosphate) pH 7.4, 150 mM NaCl (Buffer C); then 5% glycerol was added to the protein sample without lipid, or either 2 mM lyso PC/PG (50:50 mol% of 14:0 lyso PC and 14:0 lyso PG) or 1 mM β-DDM (n-Dodecyl β-D-maltoside) (*Anatrace*) were added to the samples. For characterization of the protein in buffer with no lipid, the SEC column was pre-equilibrated with Buffer C supplemented with 10% glycerol. The 10% glycerol was added, since in buffer with no glycerol the protein elution was hindered by interaction of its highly hydrophobic regions with the column beads. When studying protein in lipid environment, the column was pre-equilibrated with Buffer C supplemented by either 1 mM β-DDM or 2 mM lyso PC/PG. Then the protein (in the corresponding buffer with or without membrane mimetic) was injected in the column and eluted by running a SEC program with 0.5 ml/min flowrate. Fractions with volume of 0.5 ml were collected. In the case of protein in Buffer C with no lipid mimetics, the concentration of injected MBP-Vpu was 50 µM or 25 µM. In the case of protein in Buffer C (no glycerol) with 1 mM β-DDM, the concentration of injected MBP-Vpu was 50 µM. In the case of Buffer C (no glycerol) with 2 mM lyso PC/PG the concentration of the injected MBP-Vpu was 100 µM, 50 µM, 25 µM or 15 µM.

The fractions corresponding to protein elution peaks were combined, concentrated and analyzed by SDS-PAGE and WB as described above.

We conducted SEC on the SEC protein standards (molecular weights 12.4-200 kDa; Sigma) including β-amylase from sweet potato (200 kDa), alcohol dehydrogenase from yeast (150 kDa), albumin from bovine serum (66 kDa), carbonic anhydrase from bovine erythrocytes (29 kDa), and cytochrome c from horse heart (12.4 kDa). First, 0.4 mg of β-amylase, 0.6 mg of alcohol dehydrogenase, 1.25 mg of albumin, 0.4 mg of carbonic anhydrase, and 0.25 mg of cytochrome c were mixed in 500uL buffer C and loaded on the SEC column, pre-equilibrated with buffer C supplemented with 10% glycerol. The quantity of each protein standard was taken to produce the peaks (280 nm absorbance) of similar intensity. Later, the positions of elution peaks of MBP-Vpu in buffer, in lipid (lyso PC/PG), and in detergent (β-DDM) were referenced with the elution peaks of standard proteins.

### Preparation of samples for negative staining transmission electron microscopy (nsEM) experiments

To prepare the protein for nsEM, we exchanged first the protein buffer with buffer containing 25 mM Tris pH 7.4, 150 mM NaCl and 100 µM TCEP. Then, multiple samples of MBP-Vpu protein were prepared with and without lipids (14:0 lyso PC/PG, DHPC/DHPC). These samples can be categorized in four different types: (i) Protein in just buffer with no lipid/detergent added at protein concentrations of 0.5 µM, 1 µM, and 5 µM protein. (ii) Protein reconstituted in β-DDM at 1 µM and 5 µM protein in 1mM β-DDM, and a control of 1mM β-DDM (iii) Protein reconstituted in lyso PC/PG at 0.5 µM and 5 µM protein in 2 mM lyso PC/PG, and 5 µM protein in 1 mM lyso PC/PG plus a sample of 5 µM protein in 2 mM lyso PC/PG with 0.25 µM 5 nm GNPs and a control of 1mM lyso PC/PG; (iv) Protein reconstituted in DHPC/DHPG (06:0 PC/06:0 PG) at 0.5 µM, 5 µM and 10 µM protein in 2 mM DHPC/DHPG, 1µM protein in 0.5 mM DHPC/DHPG, 1µM protein in 1mM DHPC/DHPG, and 5 µM protein in 38 mM DHPC/DHPG plus control samples of 1 mM DHPC/DHPG and 38 mM DHPC/DHPG. The DHPC/DHPG were prepared at 50:50 mol%. Indeed, the low DHPC/DHPC concentrations at 1-2 mM are much below the critical micelle concentration of these lipids, i.e. about 15 mM, but we found these concentrations more suited for nsEM imaging. The protein was hardly visible in 38 mM DHPC/DHPG and because of this we did not continue the experiment at this lipid concentration. Each of these samples was imaged by nsEM.

### Negative staining and electron microscopy imaging

We loaded 10 µl sample onto a carbon-coated copper grid and allowed to settle for 1 min or 1 min 30 sec at room temperature (RT). Then, the solution on the grid was removed gently using filter paper and the settled on the grid protein or protein-lipid was stained by adding 10 µL of 1.5% uranyl acetate (UA), incubated for 1 min or 1 min 30 sec, and the remaining UA solution was again cleared gently using filter paper. Stained protein or protein-lipid samples were air-dried for 30 min to 1 h at RT and then used for EM imaging.

Afterward, digital micrographs were collected with a transmission electron microscope (TEM *Hitachi H-7650*) equipped with a fluorescent screen with a visual field diameter of 160 mm, 0.204 nm resolution (×20,000 final magnification), and operated at an accelerating voltage of 60 kV and 10 μA emission current. Micrographs devoid of drift and astigmatism were scanned with 1,024× 1,024 number of pixels.

The final images were visualized, contrast-adjusted and analyzed using the ImageJ software.(46)

## AUTHOR CONTRIBUTION

SM: data acquisition, data analysis and interpretation, figures, writing the manuscript; OA: data acquisition, data analysis, figures; MMI: data acquisition; BZ: data acquisition, advising; ERG: conception, design, data acquisition, data analysis and interpretation, figures, writing the manuscript, supervising, acquisition of funds. All authors participated in manuscript finalization and approved the final version of the manuscript.

## ACKNOWLEDGEMENTS

This work was supported by start-up funds from the Department of Chemistry and Biochemistry (to ERG). The College of Arts & Sciences Microscopy (CASM) is acknowledged for providing nsEM resources.

## REFERENCES

1. Khan, N., and Geiger, J. D. (2021) Role of Viral Protein U (Vpu) in HIV-1 Infection and Pathogenesis. Viruses 13

2. WHO, HIV/AIDS, https://www.who.int/data/gho/data/themes/hiv-aids, 2021.

3. Sharp, P. M., and Hahn, B. H. (2011) Origins of HIV and the AIDS pandemic. Cold Spring Harb Perspect Med 1, a006841

4. Gonzalez, M. E. (2015) Vpu Protein: The Viroporin Encoded by HIV-1. Viruses 7, 4352–4368

5. Hussain, A., Das, S. R., Tanwar, C., and Jameel, S. (2007) Oligomerization of the human immunodeficiency virus type 1 (HIV-1) Vpu protein--a genetic, biochemical and biophysical analysis. Virol J 4, 81

6. Cohen, E. A., Terwilliger, E. F., Sodroski, J. G., and Haseltine, W. A. (1988) Identification of a protein encoded by the vpu gene of HIV-1. Nature 334, 532–534

7. Strebel, K., Klimkait, T., and Martin, M. A. (1988) A novel gene of HIV-1, vpu, and its 16-kilodalton product. Science 241, 1221–1223

8. Lewinski, M. K., Jafari, M., Zhang, H., Opella, S. J., and Guatelli, J. (2015) Membrane Anchoring by a C-terminal Tryptophan Enables HIV-1 Vpu to Displace Bone Marrow Stromal Antigen 2 (BST2) from Sites of Viral Assembly. J Biol Chem 290, 10919–10933

9. Ewart, G. D., Sutherland, T., Gage, P. W., and Cox, G. B. (1996) The Vpu protein of human immunodeficiency virus type 1 forms cation-selective ion channels. J Virol 70, 7108–7115

10. Maldarelli, F., Chen, M. Y., Willey, R. L., and Strebel, K. (1993) Human immunodeficiency virus type 1 Vpu protein is an oligomeric type I integral membrane protein. J Virol 67, 5056–5061

11. Lu, J. X., Sharpe, S., Ghirlando, R., Yau, W. M., and Tycko, R. (2010) Oligomerization state and supramolecular structure of the HIV-1 Vpu protein transmembrane segment in phospholipid bilayers. Protein Sci 19, 1877–1896

12. Nieva, J. L., Madan, V., and Carrasco, L. (2012) Viroporins: structure and biological functions. Nat Rev Microbiol 10, 563–574

13. Agirre, A., Barco, A., Carrasco, L., and Nieva, J. L. (2002) Viroporin-mediated membrane permeabilization. Pore formation by nonstructural poliovirus 2B protein. J Biol Chem 277, 40434–40441

14. Fischer, W. B., and Hsu, H. J. (2011) Viral channel forming proteins - modeling the target. Biochim Biophys Acta 1808, 561–571

15. Gonzalez, M. E., and Carrasco, L. (2003) Viroporins. FEBS Lett 552, 28–34

16. Willey, R. L., Maldarelli, F., Martin, M. A., and Strebel, K. (1992) Human immunodeficiency virus type 1 Vpu protein induces rapid degradation of CD4. J Virol 66, 7193–7200

17. Strebel, K., Klimkait, T., Maldarelli, F., and Martin, M. A. (1989) Molecular and biochemical analyses of human immunodeficiency virus type 1 vpu protein. J Virol 63, 3784–3791

18. Jia, X., Weber, E., Tokarev, A., Lewinski, M., Rizk, M., Suarez, M., Guatelli, J., and Xiong, Y. (2014) Structural basis of HIV-1 Vpu-mediated BST2 antagonism via hijacking of the clathrin adaptor protein complex 1. Elife 3, e02362

19. Sato, H., Orenstein, J., Dimitrov, D., and Martin, M. (1992) Cell-to-cell spread of HIV-1 occurs within minutes and may not involve the participation of virus particles. Virology 186, 712–724

20. Klimkait, T., Strebel, K., Hoggan, M. D., Martin, M. A., and Orenstein, J. M. (1990) The human immunodeficiency virus type 1-specific protein vpu is required for efficient virus maturation and release. J Virol 64, 621–629

21. Georgieva, E. R., Borbat, P. P., Fanouraki, C., and Freed, J. H. (2020) High-yield production in E. coli and characterization of full-length functional p13II protein from human T-cell leukemia virus type 1. Protein expression and purification 173, 105659

22. Nygaard, R., Kim, J., and Mancia, F. (2020) Cryo-electron microscopy analysis of small membrane proteins. Curr Opin Struct Biol 64, 26–33

23. Liu, Y., Huynh, D. T., and Yeates, T. O. (2019) A 3.8 A resolution cryo-EM structure of a small protein bound to an imaging scaffold. Nat Commun 10, 1864

24. Georgieva, E. R., Ramlall, T. F., Borbat, P. P., Freed, J. H., and Eliezer, D. (2010) The lipid-binding domain of wild type and mutant alpha-synuclein: compactness and interconversion between the broken and extended helix forms. J Biol Chem 285, 28261–28274

25. Selmke, B., Borbat, P. P., Nickolaus, C., Varadarajan, R., Freed, J. H., and Trommer, W. E. (2018) Open and Closed Form of Maltose Binding Protein in Its Native and Molten Globule State As Studied by Electron Paramagnetic Resonance Spectroscopy. Biochemistry 57, 5507–5512

26. Hu, J., Qin, H., Gao, F. P., and Cross, T. A. (2011) A systematic assessment of mature MBP in membrane protein production: overexpression, membrane targeting and purification. Protein Expr Purif 80, 34–40

27. Lins, L., and Brasseur, R. (1995) The hydrophobic effect in protein folding. FASEB J 9, 535–540

28. Cooper, R. S., Georgieva, E. R., Borbat, P. P., Freed, J. H., and Heldwein, E. E. (2018) Structural basis for membrane anchoring and fusion regulation of the herpes simplex virus fusogen gB. Nat Struct Mol Biol 25, 416–424

29. Telmer, P. G., and Shilton, B. H. (2003) Insights into the conformational equilibria of maltose-binding protein by analysis of high affinity mutants. J Biol Chem 278, 34555–34567

30. Hutchison, J. M., Lu, Z., Li, G. C., Travis, B., Mittal, R., Deatherage, C. L., and Sanders, C. R. (2017) Dodecyl-beta-melibioside Detergent Micelles as a Medium for Membrane Proteins. Biochemistry 56, 5481–5484

31. Koehler, J., Sulistijo, E. S., Sakakura, M., Kim, H. J., Ellis, C. D., and Sanders, C. R. (2010) Lysophospholipid micelles sustain the stability and catalytic activity of diacylglycerol kinase in the absence of lipids. Biochemistry 49, 7089–7099

32. Fernandez, C., Hilty, C., Wider, G., and Wuthrich, K. (2002) Lipid-protein interactions in DHPC micelles containing the integral membrane protein OmpX investigated by NMR spectroscopy. Proc Natl Acad Sci U S A 99, 13533–13537

33. Georgieva, E. R., Borbat, P. P., Ginter, C., Freed, J. H., and Boudker, O. (2013) Conformational ensemble of the sodium-coupled aspartate transporter. Nat Struct Mol Biol 20, 215–221

34. Georgieva, E. R., Borbat, P. P., Norman, H. D., and Freed, J. H. (2015) Mechanism of influenza A M2 transmembrane domain assembly in lipid membranes. Sci Rep 5, 11757

35. Kotov, V., Bartels, K., Veith, K., Josts, I., Subhramanyam, U. K. T., Gunther, C., Labahn, J., Marlovits, T. C., Moraes, I., Tidow, H., Low, C., and Garcia-Alai, M. M. (2019) High-throughput stability screening for detergent-solubilized membrane proteins. Sci Rep 9, 10379

36. le Maire, M., Champeil, P., and Moller, J. V. (2000) Interaction of membrane proteins and lipids with solubilizing detergents. Biochim Biophys Acta 1508, 86–111

37. Georgieva, E. R., Xiao, S., Borbat, P. P., Freed, J. H., and Eliezer, D. (2014) Tau binds to lipid membrane surfaces via short amphipathic helices located in its microtubule-binding repeats. Biophys J 107, 1441–1452

38. Dube, M., Bego, M. G., Paquay, C., and Cohen, E. A. (2010) Modulation of HIV-1-host interaction: role of the Vpu accessory protein. Retrovirology 7, 114

39. Roy, N., Pacini, G., Berlioz-Torrent, C., and Janvier, K. (2014) Mechanisms underlying HIV-1 Vpu-mediated viral egress. Front Microbiol 5, 177

40. Langer, S., Hammer, C., Hopfensperger, K., Klein, L., Hotter, D., De Jesus, P. D., Herbert, K. M., Pache, L., Smith, N., van der Merwe, J. A., Chanda, S. K., Fellay, J., Kirchhoff, F., and Sauter, D. (2019) HIV-1 Vpu is a potent transcriptional suppressor of NF-kappaB-elicited antiviral immune responses. Elife 8

41. Jin, T., Chuenchor, W., Jiang, J., Cheng, J., Li, Y., Fang, K., Huang, M., Smith, P., and Xiao, T. S. (2017) Design of an expression system to enhance MBP-mediated crystallization. Sci Rep 7, 40991

42. Stangl, M., and Schneider, D. (2015) Functional competition within a membrane: Lipid recognition vs. transmembrane helix oligomerization. Biochim Biophys Acta 1848, 1886–1896

43. Duvaud, S., Gabella, C., Lisacek, F., Stockinger, H., Ioannidis, V., and Durinx, C. (2021) Expasy, the Swiss Bioinformatics Resource Portal, as designed by its users. Nucleic Acids Research 49, W216–W227

44. Omasits, U., Ahrens, C. H., Müller, S., and Wollscheid, B. (2014) Protter: interactive protein feature visualization and integration with experimental proteomic data. Bioinformatics 30, 884–886

45. Routzahn, K. M., and Waugh, D. S. (2002) Differential effects of supplementary affinity tags on the solubility of MBP fusion proteins. J Struct Funct Genomics 2, 83–92

46. Schneider, C. A., Rasband, W. S., and Eliceiri, K. W. (2012) NIH Image to ImageJ: 25 years of image analysis. Nat Methods 9, 671–675

