## Supplementary Information for "Oligomeric polymorphism of HIV-1 Vpu protein in lipid environment and in solution"

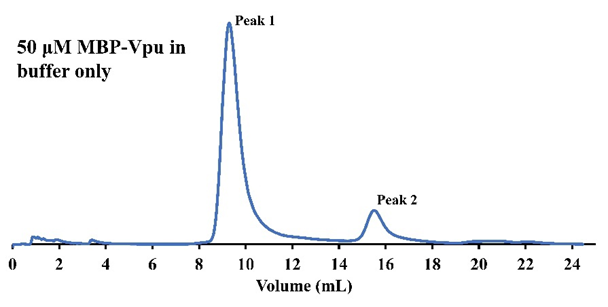

**Supporting Figure 1. SEC data for MBP-Vpu in solution**—results from second experiment identical to those in Figure 4 (main text) are shown**.** Two elution peaks (Peak 1 and Peak 2) with substantially different intensities were observed. Both Peak 1 and Peak 2 contained MBP-Vpu as confirmed by SDS-PAGE and WB (data not shown).

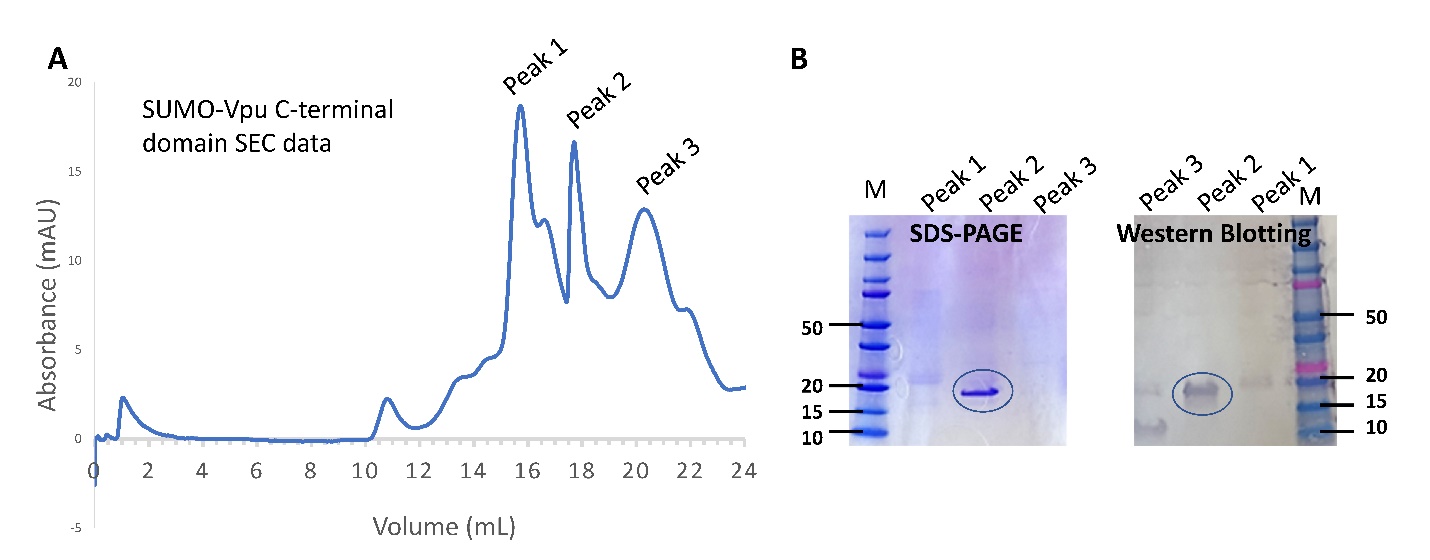

**Supporting Figure 2. Vpu C-terminal is monomeric in solution.** (A) SEC chromatogram. (B) SDS-PAGE and western blotting of SEC fractions in (A). Vpu C-terminal domain was expressed in E. coli as fusion construct with Hisx_8_-SUMO tag and purified by using consecutively Ni^2+^- and Co^2+^ - affinity chromatography, and SEC. The fractions corresponding to the three SEC elution peaks (Peak 1, Peak 2 and Peak 3 in (A)) were combined, concentrated and characterized by SDS-PAGE and western blotting. The SUMO-Vpu C-terminal protein was found in Peak 2, eluting at ca. 18 ml, which corresponds to a protein monomer of ca. 21 kDa molecular weight, based on elution of standard protein mixture (Figure 4 in the Main Text).

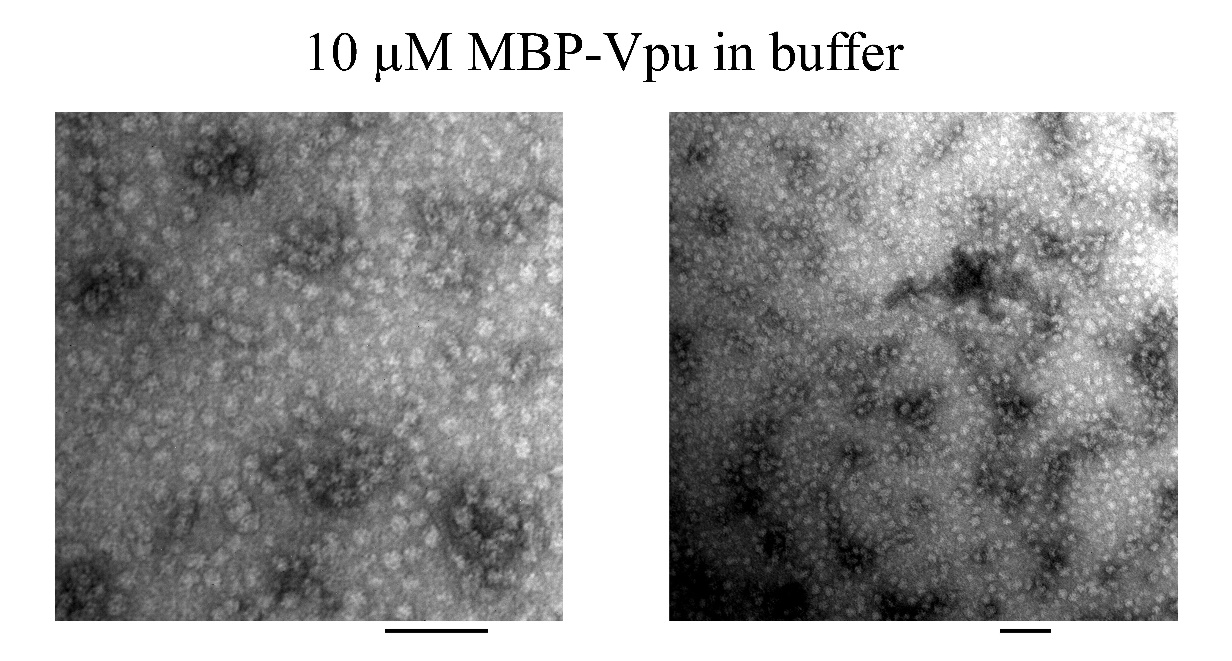

**Supporting Figure 3. Representative images of nsEM on 5µM MBP-Vpu in buffer.** The 100 nm bar is in black. Well-formed MBP-Vpu oligomers are visible.

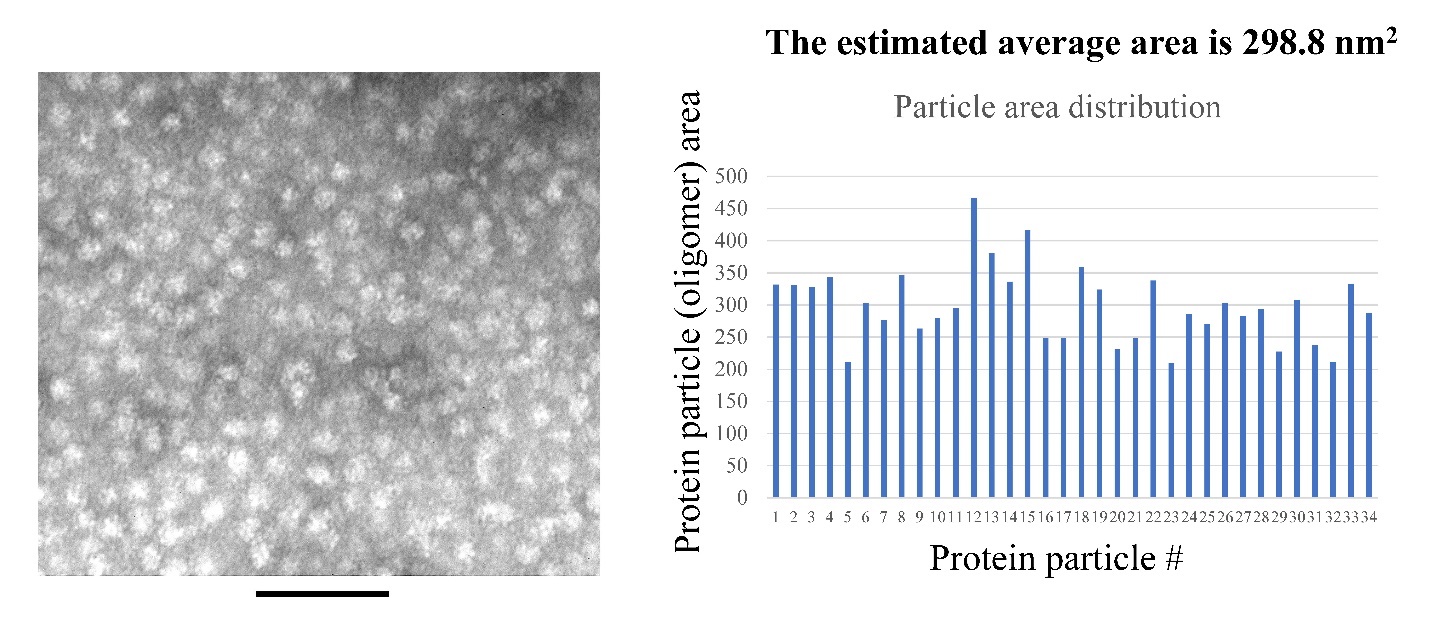

**Supporting Figure 4. Particle size distribution for MBP-Vpu oligomers in solution.** Analysis of the protein particles’ size distribution within representative area of nsEM image (left) for sample of 5 µM MBP-Vpu was conducted using the ImageJ software. A set of more than thirty particles with visually similar shape were selected manually and their area was calculated. The particle area distribution (right) is relatively narrow (except few outliers) with average particle area of 298.8 nm^2^. Ideally, this corresponds to a circle with a 9.7-9.8 nm radius and ca. 20 nm diameter.

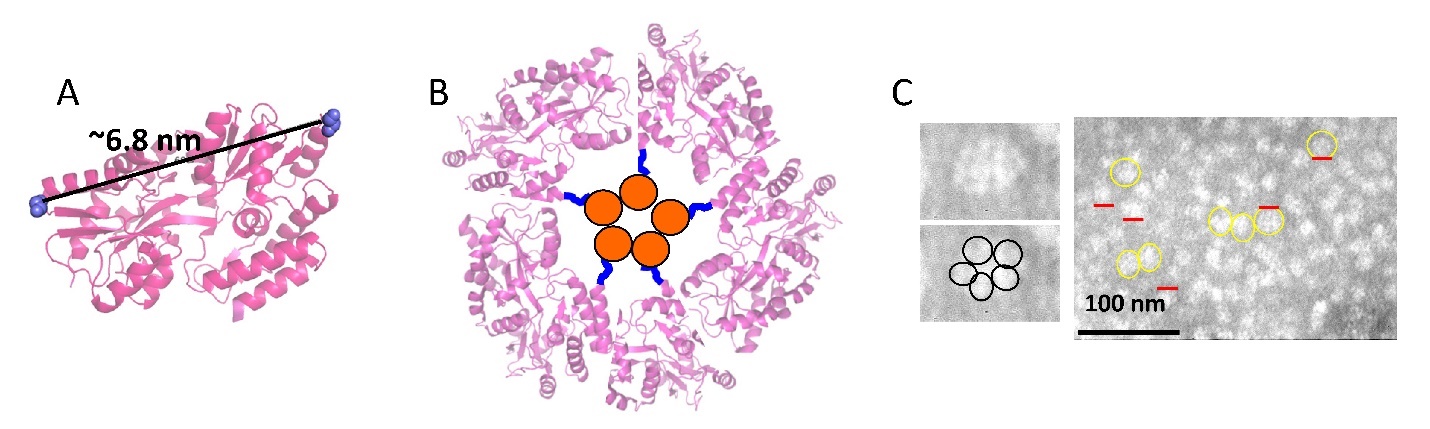

**Supporting Figure 5. Coarse model of MBP-Vpu pentamer:** (A) Crystal structure of MBP without substrate (PDB# 1PEB)—The end-to-end distance in MBP monomer is about 6.8 nm. (B) Possible arrangement of MBP-VPU monomers in the pentameric structure seen from the top—MBP moieties are as in the crystal structure and are shown in magenta, the MBP- Vpu linkers are in blue, and the TM helices of Vpu monomers are shown as orange circles. (C) The nsEM data for 5 µM MBP-Vpu in buffer (the same as in Figure 5) are shown.

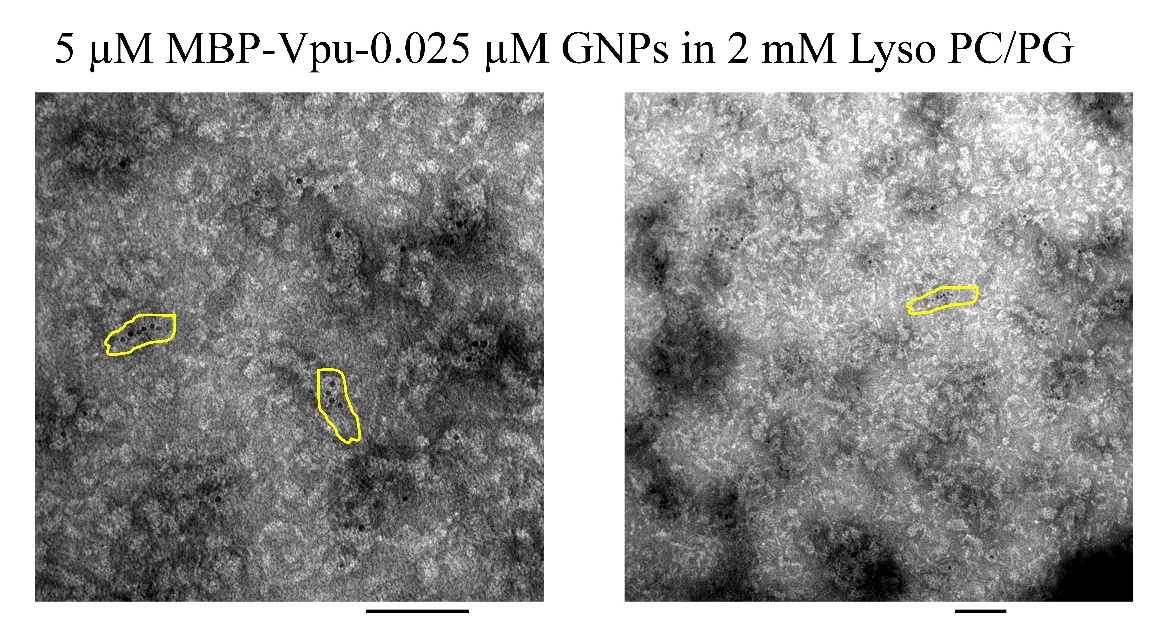

**Supporting Figure 6. Representative images of nsEM on 5µM MBP-Vpu labeled with GNPs (0.04 µM) in 2 mM lyso PC/PG.** The 100 nm bar is in black. Some of the arrays of NGP-labeled protein are enclosed in yellow. The protein was first labeled with GNPs to its His_8_-tag, and then reconstituted in lyso PC/PG. Note, not each MBP-Vpu monomer has a GNP attached, as the protein concentration was significantly higher than the concentration of GNPs.
